# A machine learning approach for detecting vicarious trial and error behaviors

**DOI:** 10.1101/2021.03.03.433785

**Authors:** Jesse T. Miles, Kevan S. Kidder, Ziheng Wang, Yiru Zhu, David H. Gire, Sheri J. Y. Mizumori

## Abstract

Vicarious trial and error behaviors (VTEs) indicate periods of indecision during decision-making, and have been proposed as a behavioral marker of deliberation. In order to understand the neural underpinnings of these putative bridges between behavior and neural dynamics, researchers need the ability to readily distinguish VTEs from non-VTEs. Here we utilize a small set of trajectory-based features and standard machine learning classifiers to identify VTEs from non-VTEs for rats performing a spatial delayed alternation task (SDA) on an elevated plus maze. We also show that previously reported features of the hippocampal field potential oscillation can be used in the same types of classifiers to separate VTEs from non-VTEs with above chance performance. However, we caution that the modest classifier success using hippocampal population dynamics is not sufficient for identifying trials where VTEs occur, and show that combining oscillation-based features with trajectory-based features degrades classifier performance compared to trajectory-based features alone. Overall, we propose a standard set of features useful for trajectory-based VTE classification and support previous suggestions that VTEs are supported by a network including, but likely extending beyond, the hippocampus.

## 1 Background and Introduction

Introduced and popularized in the 1930s, vicarious trial and error (VTE) is a well documented behavioral phenomenon where subjects vacillate between reward options before settling on their final choice (Muenzinger and Gentry, 1931; Tolman, 1938). This behavior is best understood in rats making decisions to go left or right, and as such, VTE trajectories tend to have curves that change direction at decision points. Current theories claim that subjects mentally assess possible options before making a final decision during VTEs (Redish, 2016). This makes them a valuable behavioral variable to take into account when studying decision making, particularly when investigating neural processing during decisions.

The majority of recent experiments examining the neural underpinnings of VTEs have focused on the rodent hippocampus (HPC). Bilateral electrolytic HPC lesions decrease the mean number of VTEs in a visual discrimination task, particularly during early learning (Hu and Amsel, 1995); though see (Bett et al., 2012). Similarly, bilateral ibotenic acid HPC lesions decrease the number of VTEs rats exhibit before they have located a reward in a spatial task (Bett et al., 2012). In addition to lesion studies, several electrophysiological findings link the HPC to VTEs. Dorsal HPC recordings during VTEs show serial sweeps of place cell sequences, which first trace the initial direction of the VTE before sweeping in the direction a rat ends up choosing (Johnson and Redish, 2007). Furthermore, dorsal HPC place cell recordings are more likely to represent locations of an unchosen option during VTEs than non-VTEs (Papale et al., 2016). There is also evidence that field potential oscillations recorded from dorsal HPC differ on decisions where VTEs do and do not occur. In particular, characteristics of HPC theta (4-12 Hz) oscillations, such as shape and duration, appear to be altered during VTEs, as do aspects of gamma-band (35-100 Hz) oscillations (Amemiya and Redish, 2018; Schmidt et al., 2019), but see (Dvorak et al., 2018).

Despite decades long interest and their utility as an overt marker of a putative cognitive process, VTEs have been studied by only a small number of labs. We suspect part of the reason they have not received more attention is that VTE trajectories can be highly variable, which makes it difficult to identify them algorithmically (Goss and Wischner, 1956). The Redish lab has proposed the IdPhi metric, which quantifies changes in heading angles as rats traverse choice points, for identifying VTEs (Papale et al., 2012, 2016; Redish, 2016; Amemiya and Redish, 2016, 2018; Schmidt et al., 2019; Hasz and Redish, 2020). While successful in their hands, IdPhi, admittedly, “does not provide a sharp boundary between VTE and not” (Papale et al., 2016), Supplemental Material: *Experimental Procedures*). Looking to improve upon this method of VTE identification, we demonstrate here that standard machine learning models utilizing a small number of trajectory features are able to robustly and reliably distinguish VTEs from non-VTEs in a manually scored data-set.

Additionally, we assess how the same types of classifier models perform when trained on curated features of the dorsal HPC oscillation that have been shown to differ between VTEs and non-VTEs (e.g. differences in gamma power and theta wave shape)(Amemiya and Redish, 2018; Schmidt et al., 2019). In doing so, we demonstrate that these features are indeed able to separate decision types better than would be expected by random binary classification, though with much worse performance than trajectory-based features. Furthermore, we show that providing a classifier with HPC oscillatory dynamics from when animals make choices yields better performance than oscillations from the immediately preceding delay interval, which is when information about the previous choice would need to be held in memory. We also show that a more comprehensive description of the HPC oscillation, the power spectrum, does not perform any better than the model trained on curated features. Finally, we demonstrate that combining informative trajectory- and oscillation-based features leads to a slight, but significant, decrease in classifier performance when compared to classification using trajectory features alone, leading us to conclude that the HPC oscillation does not contain information that compliments what can be extracted from the trajectories.

## 2 Results

### 2.1 Trajectory-based classification

A VTE occurs when rats vacillate between options before their final choice. Behaviorally, this manifests as a trajectory with curves or sharp angles at choice points, where reorientations occur (figure 1A). We analyzed a data set with 1137 trajectories from rats running a spatial delayed alternation (SDA) task (Baker et al., 2019; Kidder et al., 2021). Each trajectory was scored as VTE (*n* = 184) or non-VTE (*n* = 953) by at least two trained annotators. We calculated IdPhi (Papale 2012), the integrated change in heading angle, for each trajectory, as well as several other features (see 4.4 for more details). As expected, we saw statistically distinct empirical distributions for IdPhi values on trials scored as VTE compared to non-VTE (figure **1B**, *p* < 0.001, two-sample K-S test). When compared to manual scoring, however, using IdPhi did not reliably separate VTE and non-VTE trials (figure **2A**).

**Figure 1:**
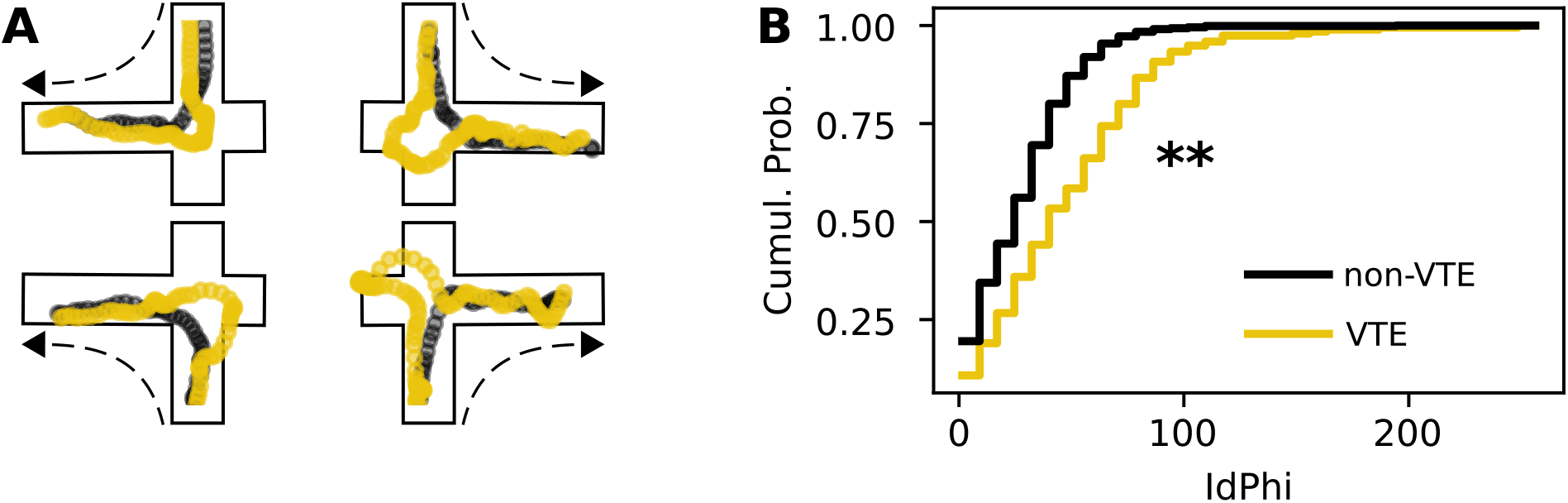
Example VTEs and IdPhi distribution. **A** - Example trajectories showing VTEs (yellow) and non-VTEs (black). Trajectories are shown on top of outlines of the decision-point, coming from each possible direction (shown by dashed arrow). **B** - Empirical cumulative distributions of IdPhi scores for VTEs (yellow) and non-VTEs (black). Note the prominent rightward shift for VTEs (p < 0.001, two-sample K-S test). *Cumul. Prob. - Cumulative probability, K-S - Kolmogorov-Smirnov*

**Figure 2:**
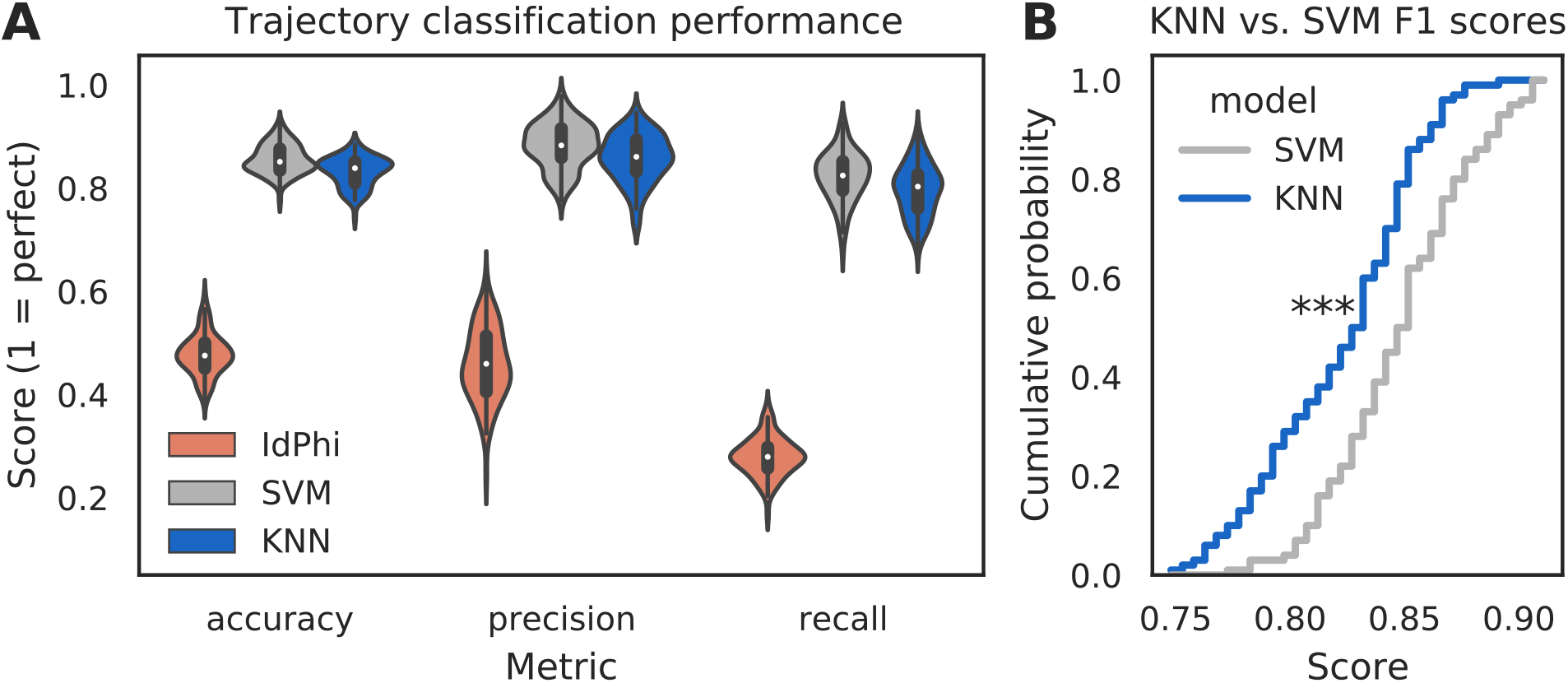
Multi-feature classifiers outperform a single metric. **A** - Classification performance on 100 random splits of manually scored data using IdPhi (orange), an SVM model (grey), and a KNN model (blue). Colored patches are kernel density estimates of the underlying distributions, with boxplots representing the same data inside the patches. **B** - Cumulative distributions of *F*_1_ scores for SVM in grey and KNN in blue. Note the rightward shift for the SVM model (*p* = 0.0001, two-sample K-S test). *SVM - support vector machine, KNN - k-nearest neighbor*

We reasoned that we could obtain more accurate and reliable VTE detection by assessing multiple aspects of the trajectory instead of just one. As such, we calculated seven trajectory-based features (see Classifier implementation for details) with the expectation that these features would allow for separation of VTEs and non-VTEs in a higher dimensional space. Like IdPhi, many of these features formed distinct empirical distributions for VTEs and non-VTEs, which suggested to us that this feature set could be used to build machine learning classifiers for algorithmic VTE detection.

Classifiers are often evaluated for their accuracy, precision, and recall scores (Malley et al., 2011; Lever et al., 2016) (see Classifier implementation for detailed descriptions). In the context of VTE identification, accuracy measures the proportion of correctly labeled trials (i.e. VTE or non-VTE), precision measures the proportion of trials labeled VTE that are actually VTEs, and recall measures the proportion of VTEs found out of the total number of VTEs present. We compared performance of two widely used machine learning models – k-nearest neighbors (KNN) and support vector machines (SVM) – to IdPhi alone in figure **2**. To generate distributions for each of these metrics, we scored 100 iterations of randomly sampled splits of data, with mutually exclusive testing and training trajectories (see Classifier implementation for further details). To ensure scores were not influenced by the fact that we had many more non-VTE trials than VTE trials, we equalized the number of VTE and non-VTE trials for each data split. Both KNN and SVM classifiers show high accuracy (*Ā_knn_* = 0.83, *Ā_svm_* = 0.86; bars above letters denote mean), precision 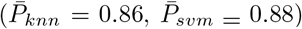, and recall 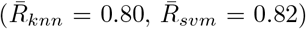 on our trajectory data. Comparing the different classifiers’ distributions of a composite precision and recall score, the *F*_1_ score, shows that the performance for the SVM classifier is generally higher (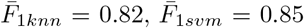; *p* = 0.0001, two-sample K-S test). Overall, these results suggest that we have defined a feature set suitable for VTE classification, that both KNN and SVM models provide more accurate, sensitive, and precise VTE classification than a single metric alone, and that the SVM model has a slight performance edge over the KNN model.

### 2.2 Oscillation-based classification

Previous research has suggested HPC involvement in decisions where VTEs occur. Early work showed that rats with bilateral HPC lesions perform less VTEs during initial learning in a visual discrimination task than rats with their hippocampi intact (Hu and Amsel, 1995). More recent research did not find differences in VTE rates for lesioned and non-lesioned animals during visual discrimination, but showed that lesioned rats exhibit fewer VTEs during early learning when performing a spatial decision-making task. In particular, lesioned rats showed fewer VTEs before finding a new reward location after it had been moved (Bett et al., 2012). Additionally, multiple studies have shown that HPC place cell activity is more likely to represent future locations during decisions involving a VTE than when no VTE occurs (Johnson and Redish, 2007), (Papale et al., 2016). Furthermore, several features of the hippocampal local field potential oscillation appear to be different when decisions are made with, as opposed to without, VTEs (Amemiya and Redish, 2018; Schmidt et al., 2019).

We tested how well features of the HPC oscillation could identify VTEs using the same approach we employed for trajectory-based VTE classification. Consistent with previous work, we found several oscillatory features with different empirical distributions for VTE and non-VTE trials (figure **3C**, bottom panel). To test whether an SVM classifier could identify VTEs above chance levels when trained with features of the HPC oscillation, we calculated classifier metric Δ scores. We compared classifier performance from two distinct behavioral epochs - one where rats actively made choices (i.e. when VTEs would occur), or during the delay interval that preceded the choice epoch. Each score shows how far above chance the classifier performed when oscillations were taken from the choice or delay epoch (figure **4A**). Chance estimates were obtained by training a classifier on oscillations from the choice epoch, but randomly labeling each trial as VTE or non-VTE. Thus, a score of zero indicates that the classifier performed the same as would be expected if randomly labeling trials. Classifier performance on HPC oscillations during choices is above the performance for classifiers trained on HPC oscillations during the delay epoch of the task (figure **4B**; 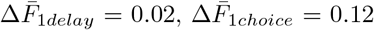; two-sample K-S test, *p* < 0.0001).

**Figure 3:**
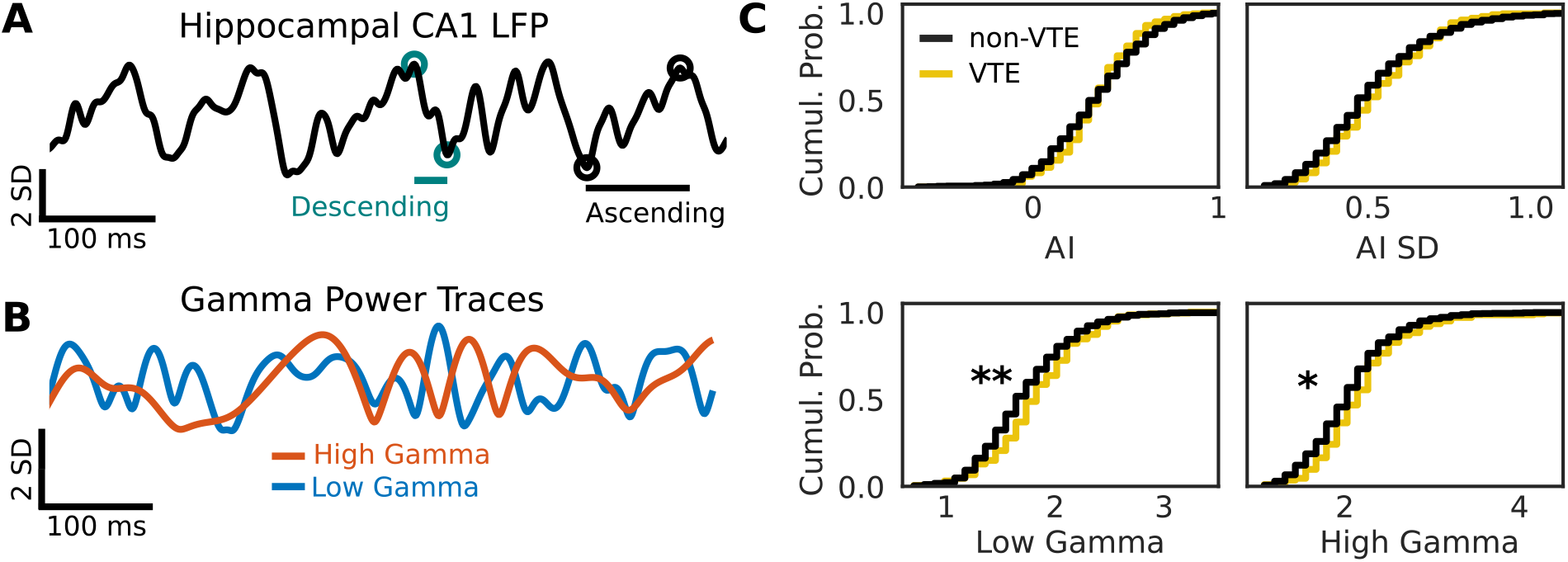
Example oscillation data and feature distributions. **A** - Sample HPC oscillation. Data are normalized (z-scored), so amplitude is measured in standard deviations (see scale bar). **B** - Normalized power timeseries for low gamma (blue) and high gamma (orange) for the oscillation shown in **A. C** - Cumulative distributions for 4 (of 12) curated features of the HPC oscillation. Asterisks denote significantly different distributions after Benjamini-Hochberg false discovery rate correction across features (Low Gamma, *p* < 0.005; High Gamma *p* < 0.05)

*AI - asymmetry index, SD - standard deviation, ms - millisecond*

**Figure 4:**
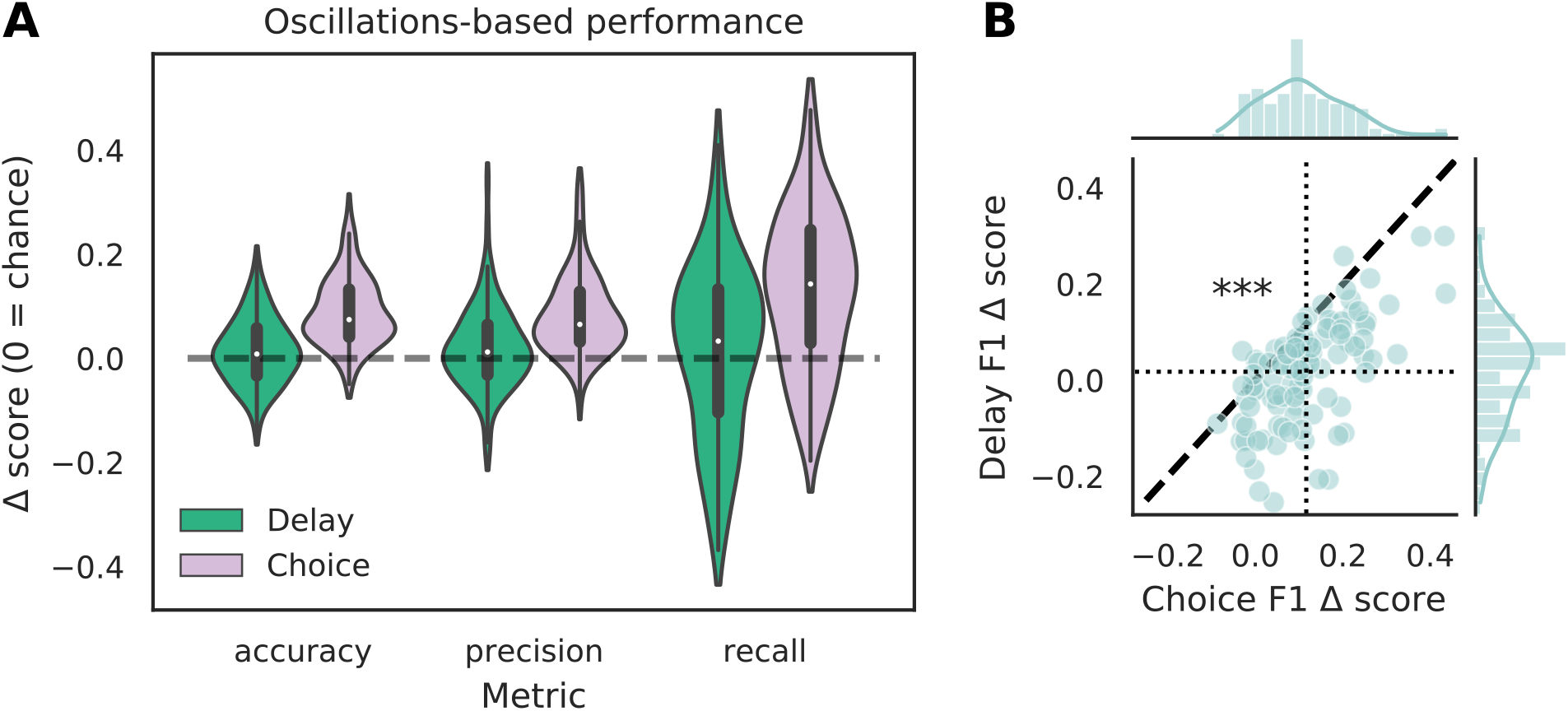
VTE classification is better for oscillation-based features from choices than delays. **A** - Classification performance when using oscillations from the delay epoch (green) compared to oscillations from the choice epoch (purple). Each Δ score is a difference between performance using data from either epoch and corresponding randomly labeled data. **B** - Scatterplots of Δ*F*_1_ score distributions for oscillations taken from the delay and choice epochs. Dotted vertical line shows the mean choice Δ*F*_1_ score, dotted horizontal line shows the mean delay Δ*F*_1_ score. Dashed diagonal line marks where equal departures from chance would occur. Individual distributions and kernel density estimates for Δ*F*_1_ scores are shown in the marginal distributions. Note skew below the diagonal, indicating significantly higher choice Δ*F*_1_ scores (*p* < 0.0001, two-sample K-S test)

Though the highly curated features used in the classifier for figure **4** have been shown to differ during VTEs and non-VTEs (Amemiya and Redish, 2018; Schmidt et al., 2019), these features are only a small subset of attributes that could describe the HPC field potential oscillation. As such, we used arguably the most common descriptor of oscillations, the power spectrum, in an attempt to increase classifier performance. We first compared average power spectral density (PSD) estimates for different frequencies, calculated for different splits of data, to identify which frequencies had significantly different average power on VTE and non-VTE decisions. Frequencies that survived false discovery rate correction (see Statistics) were used as features for an SVM classifier trained on PSD estimates. Interestingly, although these classifiers utilized a much higher dimensional feature-space (roughly seven-fold more features using PSD estimates than curated features), Δ*F*_1_ scores did not differ from those obtained with the highly curated features (figure **5B**, *p* = 0.58, two-sample K-S test).

**Figure 5:**
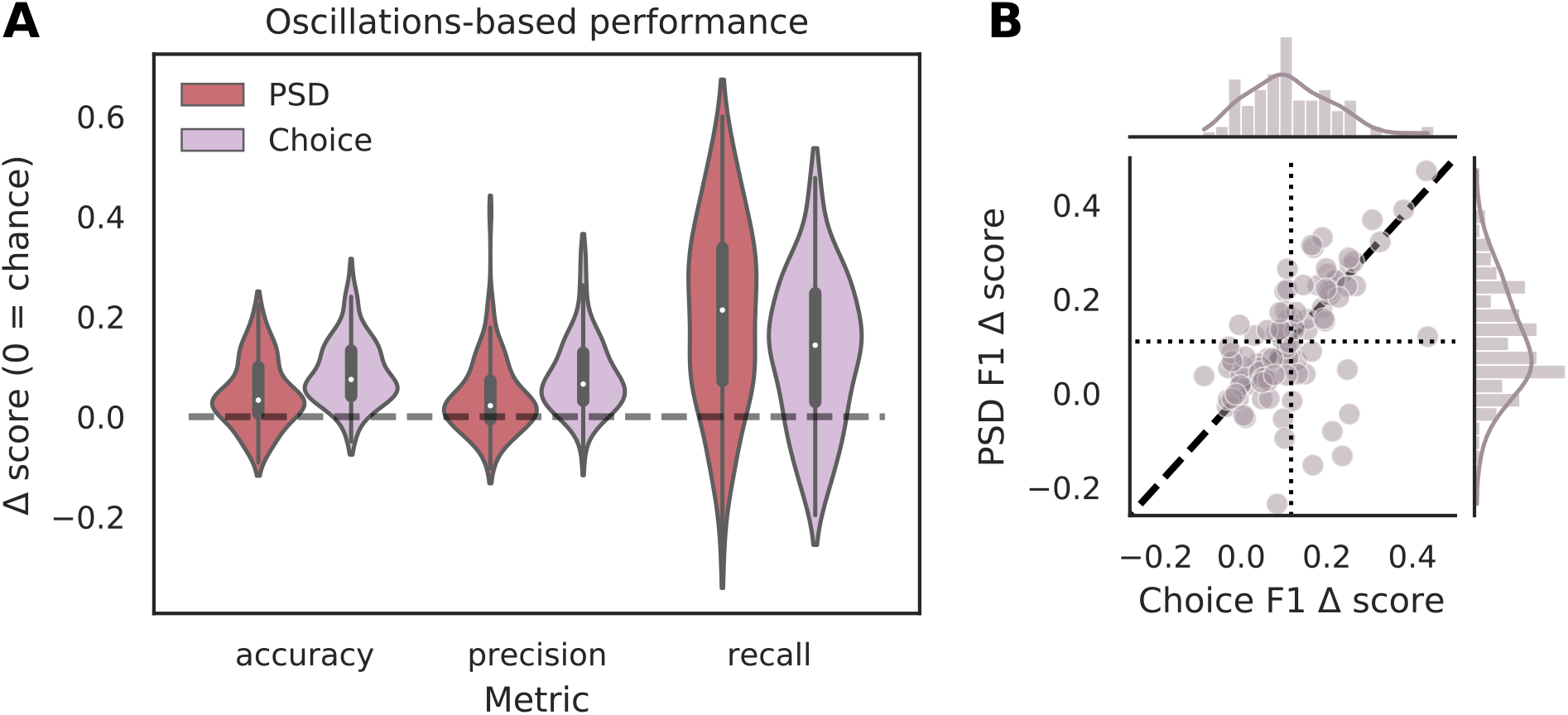
Full power spectra do not outperform highly curated oscillation features. **A** - Classification performance when using PSD estimates (dark red) compared to curated features from the choice epoch (purple). Each Δ score is measured as a difference between performance using data from either epoch and corresponding randomly labeled data. **B** - Scatterplots of Δ*F*_1_ score distributions for PSD estimates and curated features choice epochs. Dotted vertical line shows the mean choice Δ*F*_1_ score, dotted horizontal line shows the mean PSD Δ*F*_1_ score. Dashed diagonal line marks where equal departures from chance would occur. Individual distributions and kernel density estimates for Δ*F*_1_ scores are shown in the marginal plots. Δ*F*_1_ scores are not significantly different for SVMs trained on PSD-based and curated feature sets (*p* = 0.58, two-sample K-S test). *PSD - power spectral density*

It is possible that features of the HPC oscillation contain information about VTE occurrence that compliments the information contained in trajectory data. In other words, VTEs that are difficult to classify based on trajectories alone may have accompanying HPC oscillatory dynamics that, when combined with the trajectory features, lead to improved VTE classification. To examine this possibility, we trained an SVM classifier on combined trajectory and curated oscillation-based features that had significantly different distributions on VTE and non-VTE trials. Interestingly, combining feature sets (slightly) decreased performance when compared to trajectory features alone (figure **6**). Thus, we conclude that, although features of the HPC oscillation can be used to some extent for classifying VTEs, these features do not contain novel or complementary information beyond what can be extracted from the trajectories.

**Figure 6:**
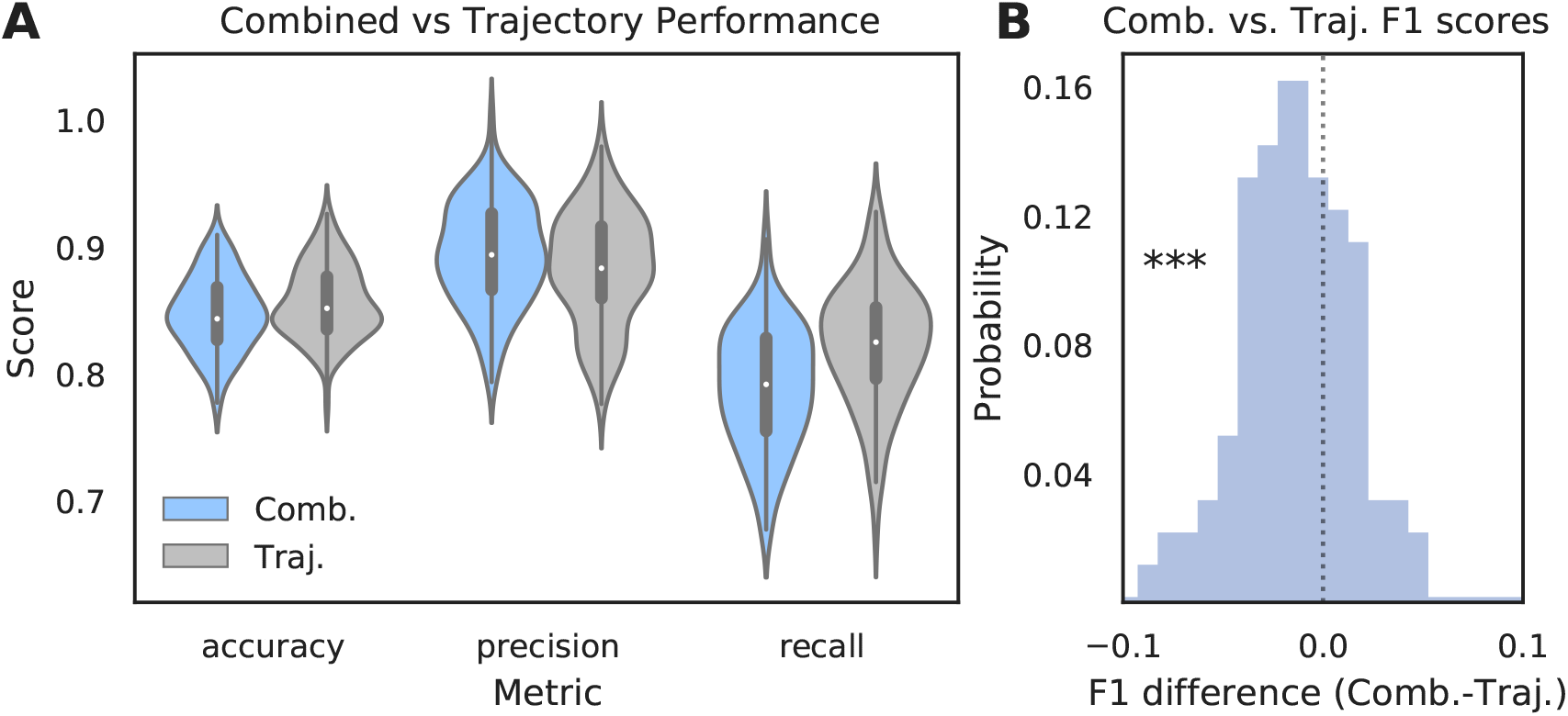
Adding oscillations-based features slightly degrades classifier performance. **A** - Comparison of SVM classifier performance using both oscillation- and trajectory-based features (purple) or trajectory features alone (grey, same as figure **2 A** - SVM). **B** - Distribution of F1 Δ scores for different data splits. Values below zero indicate worse performance using combined features. Scores are significantly shifted below zero (p < 0.0001, two-sample K-S test) *Comb. - Combined, Traj. - Trajectory alone*

## 3 Discussion

The purpose of this study was to improve upon current methods of VTE identification and build on our understanding of hippocampal involvement during VTEs. We show that VTE behavior can be robustly and reliably separated from non-VTE behavior using a small set of trajectory-based features. Additionally, we show that classifiers trained on features of the dorsal HPC field potential oscillation separate VTEs from non-VTEs more than would be expected by chance, supporting previous research linking the HPC to VTEs. Moreover, we show that when oscillations are taken from the delay epoch that precedes the choice epoch, oscillation-based features no longer enable above chance VTE classification, which suggests a brief temporal window underlying HPC involvement in VTE processing. We also caution, however, that despite above chance VTE identification using oscillation-based features, our results also clearly show that population level HPC dynamics are not sufficient for robust VTE identification (figures **4** and **5**). In particular, we demonstrate that combining neural features and trajectory features leads to slightly, but significantly, worse performance than using trajectory features alone, suggesting that the HPC oscillation-based features do not contain information that compliments or adds to the behavioral features.

Not only do curated HPC oscillation-based features show only modest VTE classification, we show that full power spectra do not improve performance, despite using roughly seven-fold more features. We see this as further evidence that population level HPC dynamics are not sufficient to identify VTEs, but acknowledge that taking information from place cell sequences into account might bolster performance, given the reports of sequences tracing out upcoming VTEs before the behavior occurs (Johnson and Redish, 2007). In our opinion, however, it is more likely that examining HPC interactions with other areas, such as the medial prefrontal cortex (mPFC). (Brown et al., 2016; Voss and Cohen, 2017; Schmidt et al., 2019; Hasz and Redish, 2020; Kidder et al., 2021), would be more fruitful for improving our ability to classify VTEs based on neural activity. Schmidt *et al.* have shown that rats perform fewer VTEs when faced with difficult decisions if their mPFC has been inhibited chemogenetically. Furthermore, the window in which HPC oscillations are best able to identify VTEs is during choices, which is when brief increases in theta coherence between the dorsal HPC and mPFC occurs (Jones and Wilson, 2005; Benchenane et al., 2010), suggestive of cross-regional communication (Fries, 2005, 2015). Finally, experiments using optogenetics to perturb the mPFC in a taskepoch-specific manner during the SDA task showed that stimulation decreased the proportion of VTEs rats engaged in, with a trend toward choice epoch mPFC disruption having a greater effect than stimulation in other epochs (Kidder et al., 2021).

Methodologically, we find comparing classification performance between behavior and neural activity an intuitive way to understand how well the activity under scrutiny relates to the behavior in question. When behavior classification is good, that level of performance can often be thought of as an upper bound for assessing how well neural activity describes the behavior, while randomly labeled classifiers can set the lower performance bound. This may provide a more nuanced picture of how well neural activity describes a behavior than hypothesis testing alone. For example, while we and others show multiple features of the HPC oscillation form distinct empirical distributions for VTEs and non-VTEs, the fact that classifier performance using these features does not meet classification performance of the behavior itself suggests that these features only provide a partial description about the neural substrate of the behavior. Additionally, feature-based classification allows for very flexible control of what parameters - behavioral or neural - one wishes to examine, as well as the size of the parameter space one would like to search. Moreover, as demonstrated by comparing HPC power spectra with curated oscillation features, feature vectors can be arbitrarily sized with surprisingly little influence on classifier performance, as long as the classifier is constructed to protect against overfitting (e.g. with proper hyper-parameter selection and cross-validation). For these reasons, we see this framework as extremely flexible in terms of feature selection and use, as well as an intuitive way of gauging how well neural activity measurements describe behavior.

Altogether, our results expand previous efforts to algorithmically identify VTEs based on trajectories, improving our ability to detect these important variants of decision-making behavior. In addition, we provide support for hypotheses that position the hippocampus as one element in what is likely a broader network of interacting neural structures that support VTEs. We believe future decision-making research will benefit from tracking VTEs and VTE-like behaviors, such as saccades and head movements in humans and non-human primates (Voss and Cohen, 2017; Santos-Pata and Verschure, 2018) and hope our classification scheme enables more wide-spread VTE analysis. Additionally, we encourage future VTE research to expand beyond the HPC and further our understanding of the neural system(s) involved in this decision-making behavior.

## 4 Methods

### 4.1 Behavioral task

Food restricted (85% of body weight) Long Evans rats (n = 9, Charles River Laboratories) were trained on a previously described spatial delayed alternation (SDA) task (Baker et al., 2019), (Kidder et al., 2021). Briefly, sessions were run on an elevated plus maze (black plexiglass arms, 58 cm long x 5.5 cm wide, elevated 80 cm from floor), with moveable arms and reward feeders controlled by custom LabView 2016 software (National Instruments, Austin, TX, USA). Each trial consisted of a rat leaving its starting location in a randomly chosen “north” or “south” arm, then navigating to an “east” or “west” arm for a 45 mg sucrose pellet reward (TestDiet, Richmond, IN, USA). Rewards were delivered when rats alternated from their previously chosen arm (i.e. if they selected the “east” arm on trial *n* − 1 then they had to select the “west” arm for reward on trial n). After making a choice, rats had the opportunity for reward consumption (if correct) before they returned to the assigned start arm and entered into a 10 second delay period before the next trial began. Based on this structure, we divided the task into three epochs - choice, return, and delay.

### 4.2 Microdrive implantation

Micro-drive bodies were 3-D printed (Form 2 Printer; Formlabs, Sommerville, MA) to contain between 8 and 16 gold plated tetrodes (nichrome; SANDVIK, Sandviken, Sweden), which were implanted unilaterally into the CA1 region of HPC (AP: −3.0, M/L: ±2.0mm, D/V: −1.8mm). A subset of animals (3) had two optic fibers that were implanted bilaterally into the mPFC and used for additional experiments (Kidder et al., 2021), and the remainder (6) also had tetrodes implanted into the ipsilateral lateral habenula for additional experiments, but all animals ran the same behavioral task, and data used for this study were from before any optogenetic stimulation was ever delivered. Tetrodes were connected to a 64-channel Open Ephys electrode interface board (EIB) (open-ephys.org). To eliminate external noise, drive bodies were shelled in plastic tubes lined with aluminum foil coated in a super-conductive nickel spray. One ground wire connected the shell with the EIB and, during surgery, another ground wire was implanted near the cerebellum just inside the skull. After surgery, rats were allowed to recover for approximately seven days before entering into testing, and HPC tetrodes were lowered over the course of several days until at least one tetrode showed oscillations consistent with the CA1 fissure (high-amplitude, asymmetric theta oscillations).

### 4.3 Data Acquisition

#### Behavior tracking

Two LEDs were attached to either the rat’s microdrive or the tethers plugged into the microdrive’s head-stage before recordings. Rat locations were determined by subtracting a background image taken at the beginning of the session from each frame. Pixels containing the LEDs showed an above threshold difference in brightness, which allowed us to determine rat head locations in each frame. Camera frames were recorded at approximately 35 Hz using a SONY USB web camera (Sony Corporation, Minato, Tokyo). Frames were time-stamped with a millisecond timer run by LabView and sent to the Open Ephys acquisition software (open-ephys.org) for later alignment of electrophysiological and position information.

#### Electrophysiology

Electrophysiological data were sampled at 30 kHz using Intan headstages (RHD2132; Intan Technologies, Los Angeles, California) connected to the Open-Ephys EIB. Digitized signals were sent via daisy chained SPI cables through a motorized commutator that prevented tether twisting (AlphaComm-I; Alpha Omega Co., Alpharetta, GA) and into an Open-Ephys acquisition board (open-ephys.org). All further processing and filtering was done offline using custom MATLAB scripts (see section 4.5 for more details).

### 4.4 Classifier features - trajectories

We calculated 7 features of choice epoch trajectories - the standard deviation (SD) of the x-position (*x_σ_*), the SD of the y-position (*y_σ_*), the trial’s integrated change in heading angle (*IdPhi*), the trial duration (*dur*), how well the trial was fit by a sixth degree polynomial (*r*^2^), and the number of Fourier coefficients needed to describe the fit of the polynomial (*n_coef_*). Both *x_σ_* and *y_σ_* were calculated using the std method from `Python’s numpy` package for the x and y position vectors of the rat’s trajectory on a given trial. The *IdPhi* score for a trial was defined as:

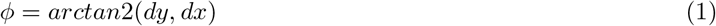

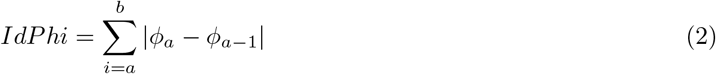

where *arctan2* is the 2-argument arctangent function and *dx* and *dy* are changes in the trajectory’s x and y position, respectively. We set the *IdPhi* threshold value, above which something was assigned as a VTE, by iterating through values from the 50th to the 80th percentile and choosing the value that maximized classification accuracy. The *dur* feature measures the duration a rat was within an experimenter defined choice point on the maze. The *r*^2^ value was determined using a two-step process. First, optimal coefficients for each of the terms in the polynomial were calculated using the `curve_fit` method of the `scipy.optimize` package with the vector of x positions as the independent variable and the vector of y positions as the dependent variable. From here, we used the optimized outputs as inputs to a generic sixth degree polynomial function, calculated the error sum of squares between the observed y values and modeled outputs (*SSE*, see equation 3), and calculated the total sum of squares (SST, see equation 4). The calculation of the *r*^2^ value is shown in equation 5.

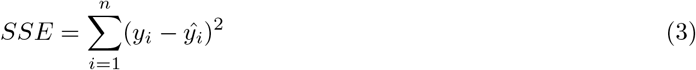

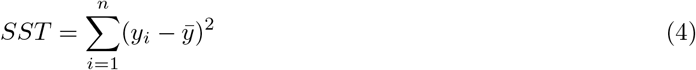

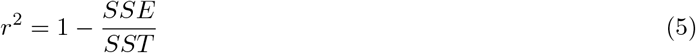

In (3), 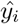 is the estimated y position at the *i*-th location in the trajectory, and 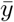 is the mean y position of the trajectory in (4). We noticed that plotting the polynomial estimates with poor fits (which were mainly VTEs) created a trajectory that looked similar to a damped oscillation, so we devised the *n_coef_* feature - which is the number of Fourier coefficients needed to describe the polynomial fit estimate - to capture this oscillatory character. Intuitively, higher values of *n_coef_* were expected to correlate with instances of VTE.

### 4.5 Classifier features - curated oscillation features

To quantify features of the HPC CA1 oscillation, we down-sampled our data by a factor of 30, going from a 30kHz sampling rate to a 1kHz sampling rate, and z-scored the downsampled timeseries to put amplitude in units of standard deviations. Based on previous work (Amemiya and Redish, 2018; Schmidt et al., 2019), we were interested to see if we could use features of the HPC CA1 oscillation to classify VTE vs non-VTE trials. We used 7 features - the asymmetry index (*AI*) of the wide-band theta oscillation, average ascending (*asc*) and descending (desc) durations of the wide-band theta oscillation, the average (normalized) low and high gamma powers (*LG* and *HG*, respectively), the cycle-averaged gamma ratio (*GR*), and the average duration of a theta cycle. Each trial had multiple measurements of each value, so we also used the SD of these measurements as a feature for all but the *asc* and *desc* features, giving a 12-variable feature vector for each trial.

Previous reports have demonstrated asymmetric theta oscillations in different layers of the HPC (Buzsáki et al., 1985, 1986; Buzsáki, 2002), so we used a low-pass filtered signal with the cutoff frequency at 80 Hz to identify peaks and troughs of the theta oscillation as well as the ascending duration, descending duration, and total duration of each theta cycle (Belluscio et al., 2012) (see figure **3A** for an example). The *AI* is defined as:

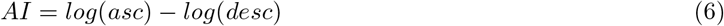

such that cycles with longer ascending than descending durations will give positive values, cycles with equal ascending and descending durations will equal 0, and cycles with shorter ascending than descending durations will give negative values. Because different HPC recording locations can have differently shaped theta oscillations (Buzsaki et al., 1985, 1986; Buzsaki, 2002), we ensured that all days used for analysis had *AI* distributions that were skewed in the same direction.

To estimate gamma powers, first we bandpass filtered our downsampled timeseries between 35-55 Hz for low gamma and 61 - 100 Hz for high gamma using third order, zero-lag Butterworth filters. These values were then z-scored, putting units of amplitude into standard deviations. The power in a gamma-band timeseries, g(t), was estimated using:

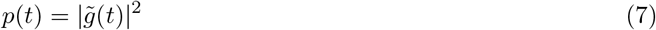

where 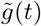 denotes the Hilbert transform of *g(t)*. We then used these power estimates to calculate cycle-by-cycle *GRs*. For a given cycle, the gamma ratio was defined as:

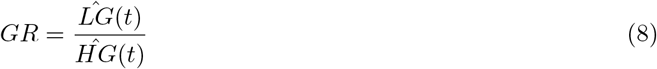

and the gamma ratio for the entire trial was the average of these cycle-by-cycle values.

### 4.6 Classifier features - power spectral density

In addition to pre-defined oscillation bands and bandpass filtering signals, we performed the same classifier-based analysis of neural data using PSD estimates as features instead of the curated oscillation features. For this, we used MATLAB’s `periodogram` function (version 2018 B; MathWorks, Nattick, MA), with a Hamming window over the duration of the signal, a frequency resolution of 1 Hz, and a range of 1 Hz to 100 Hz. To maintain consistency with curated oscillation features, we use the z-transformed HPC oscillation. PSD estimates are kept as original values, as opposed to the common decibel conversion.

### 4.7 Classifier implementation

#### Classifier models

We used the `scikit_learn` library from `Python` to create and test k-nearest neighbor (KNN) and support vector machine (SVM) models. All instances of the KNN model used 5 neighbors for classification, though results for 3-10 neighbors did not lead to different conclusions. All instances of the SVM model used a radial basis function (RBF) kernel for assessing distance/similarity. A *γ* parameter dictates the width and shape of the RBF, with lower values giving wider kernel functions and higher values giving narrower kernel functions. We chose to search values between 0.005 and 10 for *γ*. Another parameter, the C parameter, controls the trade-off between the size of the decision function margin and classification accuracy, which can be thought of as a way to control overfitting the decision function. Low values of C favor a larger margin, high values of C favor a more complex decision function. We tested a range of C values from 0.01 to 10. Data used to train the model were standardized and scaled. Testing data given to the model were transformed based on the scalings calculated for the training data (see *Cross Validation* for more details on how data were used for classifier training and testing).

#### Evaluating classifiers

We used several standard metrics for assessing classifier performance (Lever et al., 2016), all of which describe different combinations and/or weightings of true positives (*TP*), true negatives (*TN*), false positives (*FP*), and false negatives (*FN*). For VTE identification, a *TP* is trial correctly classified as a VTE, a *TN* is a trial correctly classified as a non-VTE, a *FP* is a trial incorrectly classified as a VTE, and a *FN* is a trial incorrectly classified as a non-VTE. Accuracy measures the number of trials assigned to the correct class (VTE or non-VTE) out of the total number of trials, and is defined as:

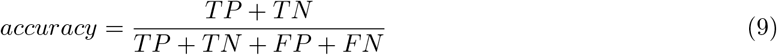

such that accuracy equals 1 if every trial, VTE and non-VTE, is correctly classified, and 0 if no trials are correctly classified. Precision measures the number of correctly classified VTEs out of the total number of trials classified as a VTE, i.e.:

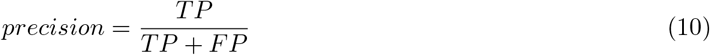

meaning precision takes a value of 1 if all of the trials classified as a VTE are in fact VTEs, even if it does not identify all VTEs in the dataset. As a complimentary metric, recall takes *FN* into account:

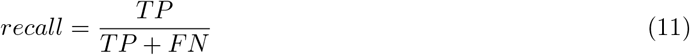

and is thus a measure of how many VTEs were correctly classified out of the total number of VTEs in the dataset. For a binary classifier with equal numbers of each class, chance performance for each metric would be 0.5 on average.

It is often helpful to compare accuracy, precision, and recall, individually but a composite measure that combines precision and recall scores is the F-score. For this paper we summarize classifier performance with the F1-score (*F*1) which is defined as follows:

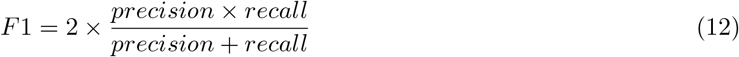

#### Cross validation

To ensure our classifiers were generalizable and performance was not biased by a particular ordering of our dataset, we performed cross-validation on distinct test/train splits of the dataset. For each evaluation, we used 67% of data for supervised training, and used the remainder for testing performance. For reproducibility, and to make comparisons across classifier models and feature modalities, we created a (seeded) matrix of randomly shuffled trials where each column contained a distinct ordering of trial values to use for one split of model training and testing (Liu, X.-Y. et al., 2009). For a given assessment, we used 100 distinct splits of testing and training data, giving a matrix with 100 columns. Since VTEs occur on roughly 20% of trials, every VTE in the dataset was present in each column, and a randomly drawn, equal number of non-VTEs made up the rest of the column, meaning each distinct split used the same VTE trials, but was allowed to contain different non-VTE trials (Liu, X.-Y. et al., 2009). This same matrix was used any time we evaluated classifier performance, meaning all evaluations were done using the exact same 100 iterations of test/train splits. Put simply, we assessed performance with 100 iterations of randomly selected trials constituting each test/train split, but ensured that assessments for different classifier models and feature modalities were performed on the exact same data. All hypotheses were tested using seed values of 1 through 5 with the `numpy.random.default_rng(seed)` method, and all conclusions were the same with each value of seed. The figures in this paper were generated using `seed = 5`.

### 4.8 Dataset curation

Training and assessing performance of the supervised classifier required manual VTE scoring to assign labels to trials. Because it is difficult to define an exact set of criteria for scoring a VTE (hence the need for a classifier), we instead chose to have at least two raters score each trial, and used their consensus to determine the label. All raters first trained together on a subset of trials to develop “implicit criteria” for scoring.

We also excluded trials based on several criteria of the hippocampal oscillations. First, we checked that the overall central tendency of the *AI* distribution was positive for a given session. Note that other studies have reported generally negative *AIs* (Amemiya and Redish, 2018; Schmidt et al., 2019). We suspect this is due to systematic shifts in theta shape characteristics across the different hippocampal axes (Buzsáki et al., 1985, 1986; Buzsáki, 2002). We also excluded trials where a 4 SD noise threshold, calculated based on the SD of the entire timeseries, was exceeded. If any session had more than 25% of its trials excluded, we did not use any of the data from that session.

### 4.9 Statistics

We performed two-sample, two-tailed Kolmogorov-Smirnov (K-S) tests to evaluate whether empirical distributions are likely drawn from the same underlying population distribution. To test whether a distribution of differences is centered at zero (i.e. to test for differences between paired groups), we performed one-sample, two-tailed Wilcoxon signed-rank tests. To assess which features exhibit statistically distinct empirical distributions when testing a number of features, we follow K-S testing with Benjamini-Hochberg (BH) false discovery rate correction to adjust *p*-values. Criteria for significance is set at *p* = 0.05 for all tests and corrections.

## Author Contributions

JTM, KSK, DHG, and SJYM contributed to conception and design of the study. JTM, KSK, ZW, and YZ annotated data. JTM analyzed the data. JTM wrote the manuscript. All authors contributed to manuscript revision, read, and approved the submitted version.

## Acknowledgments

The authors thank Victoria Hones and Yingxue Rao for assisting with behavioral training and neural recordings. We thank Victoria Hones, Brian Jackson, Mohammad Tariq, and Philip Baker for comments on the manuscript.

